# A Sensory-Motor Theory of the Neocortex based on Active Predictive Coding

**DOI:** 10.1101/2022.12.30.522267

**Authors:** Rajesh P. N. Rao

## Abstract

We propose that the neocortex implements active predictive coding (APC), a form of predictive coding that incorporates hierarchical dynamics and actions. In this model, each neocortical area estimates both sensory states and actions, and the cortex as whole learns to predict the sensory consequences of actions at multiple hierarchical levels. “Higher” cortical areas maintain more abstract representations at larger spatiotemporal scales compared to “lower” areas. Feedback from higher areas modulate the dynamics of both state and action networks in lower areas. This allows the cortical network to model the complex dynamics and physics of the world in terms of simpler compositional elements (state transition functions). Simultaneously, current higher level goals invoke sequences of lower level sub-goals and actions, allowing the network to solve complex planning problems by composing simpler solutions. Planning (“system 2” thinking) in turns allows the network to learn, over time, perception-to-action mappings (policies; “system 1” thinking) at multiple abstraction levels. We provide examples from simulations illustrating how the same APC architecture can solve problems that, at first blush, seem very different from each other: (1) how do we recognize an object and its parts using eye movements? (2) why does perception seem stable despite eye movements? (3) how do we learn compositional representations, e.g., part-whole hierarchies, and nested reference frames for equivariant vision? (4) how do we model the “physics” of a complex environment by decomposing it into simpler components? (5) how do we plan actions in a complex domain to achieve a goal by composing sequences of sub-goals and simpler actions? and (6) how do we form episodic memories of sensory-motor experiences? We propose a mapping of the APC network to the laminar architecture of the cortex and suggest possible roles for cortico-cortical, cortico-thalamic, cortico-hippocampal and cortico-subcortical pathways.

## 1 Introduction

In 1999, the author (with Dana Ballard) proposed a theory of visual processing based on hierarchical predictive coding [1]. This theory postulated that the neocortex implements a hierarchical generative model of images such that feedback connections between cortical areas carry predictions of neural activity from a higher to a lower area while feedforward connections convey the prediction errors, allowing the higher area to correct its predictions (see also [2]).

While this theory has been the subject of increasing attention in recent years (for a review, see [3]), it fails to acknowledge a fundamental aspect of perception, namely, that perception is action-based: we move our eyes about three times a second to recognize objects in a scene, orient our heads to localize sounds in our environment, and use our fingers to identify objects by touch. Away from experimentally-imposed constraints in the laboratory, perception, in its natural state, can best be viewed as an action-based hypothesis testing process.

Such a view harmonizes well with the observation that all areas of the neocortex (henceforth, the “cortex”), including the areas traditionally labeled sensory cortices, send outputs to subcortical motor regions (see [4] and references therein). Indeed, Vernon Mountcastle, in his prescient article in 1978 [5], put forth the idea that a single unifying computational principle might be operating across the entire cortex, proposing the “cortical column” as a modular information processing unit of the cortex (related ideas can be found in [6, 7]).

If there is indeed a single computational principle operating across cortex, how could it explain capabilities as diverse as (a) learning to recognize an object from foveal images or touch through eye or finger movements, (b) solving a complex spatial navigation task using simpler movement sequences, (c) understanding abstract concepts such as a family tree?

In this article, we explore active predictive coding (APC), a new theory of the cortex that acknowledges the fundamental role of hierarchical sensory-motor models in cortical function. Specifically, we take seriously the suggestions of Mountcastle and others [4, 5, 7] and propose a canonical cortical module as consisting of a *sensory state prediction network* and an *action prediction network*, both implemented within a cortical column. The two networks are tightly coupled to each other, with state predictions in superficial layers in a column feeding to and receiving feedback from deeper layers in the column. Feedback from higher cortical areas modulates the dynamics of both networks, leading to representations that operate at multiple levels of sensory and motor abstraction, as observed in cortical hierarchies implicated in perception and action [8, 9]. The author’s team has previously presented the computational elements of APC to AI audiences [10–12]. Here we explore the application of APC to understanding cortical function.

## 2 Active Predictive Coding

### 2.1 Neuroanatomical and Physiological Motivation

The theory of active predictive coding is motivated by the growing body of experimental evidence that (1) almost all cortical areas, even primary sensory ones, are influenced by actions, and (2) the axonal outputs of layer 5 neurons in almost all cortical areas target subcortical motor centers [4, 13]. This has lead to the suggestion that such layer 5 outputs across the cortex can be regarded as “motor” outputs [4]. Indeed, even in primary visual cortex (V1), outputs from layer 5 neurons target the superior colliculus [14], which is involved in eye movements. Similarly, layer 5 neurons in primary auditory cortex (A1) send outputs to the inferior colliculus [15] which is involved in orienting and defensive motor behaviors [16] while layer 5 neurons in the primary somatosensory cortex (S1) send outputs to the spinal cord [17] which controls body movements.

Figure 1A shows the laminar structure of a typical cortical column and its connectivity (based on [4, 18, 19]). Inputs from a sensory region or a lower cortical area enter layer 4 whose outputs are then conveyed to the superficial layers 2/3 neurons. These neurons in turn send their axons to the deeper layers, predominantly targeting layer 5 neurons, which are considered the “output” neurons of the cortical column. One class of layer 5 neurons (with thick tufted apical dendrites and firing in bursts) send their axons to subcortical motor centers such as the superior colliculus and other parts of the brainstem [14, 19]. These layer 5 neurons have thus been called “motor output” neurons and occur across cortical areas, including primary sensory cortices [4]. Other layer 5 neurons, which do not fire in bursts and have slender apical dendrites, project to the striatum and other cortical regions [14, 19]. There is also a significant axonal projection from layer 5 back to layer 2/3, signifying recurrent feedback within a cortical column. There are additional projections from Layer 5 to Layer 6, which in turn sends outputs to the part of the thalamus that sending inputs to Layer 4. These outputs from Layer 6 also send collaterals to the inhibitory neurons of the thalamic reticular nucleus (TRN) which can inhibit thalamic neurons.

**Figure 1:**
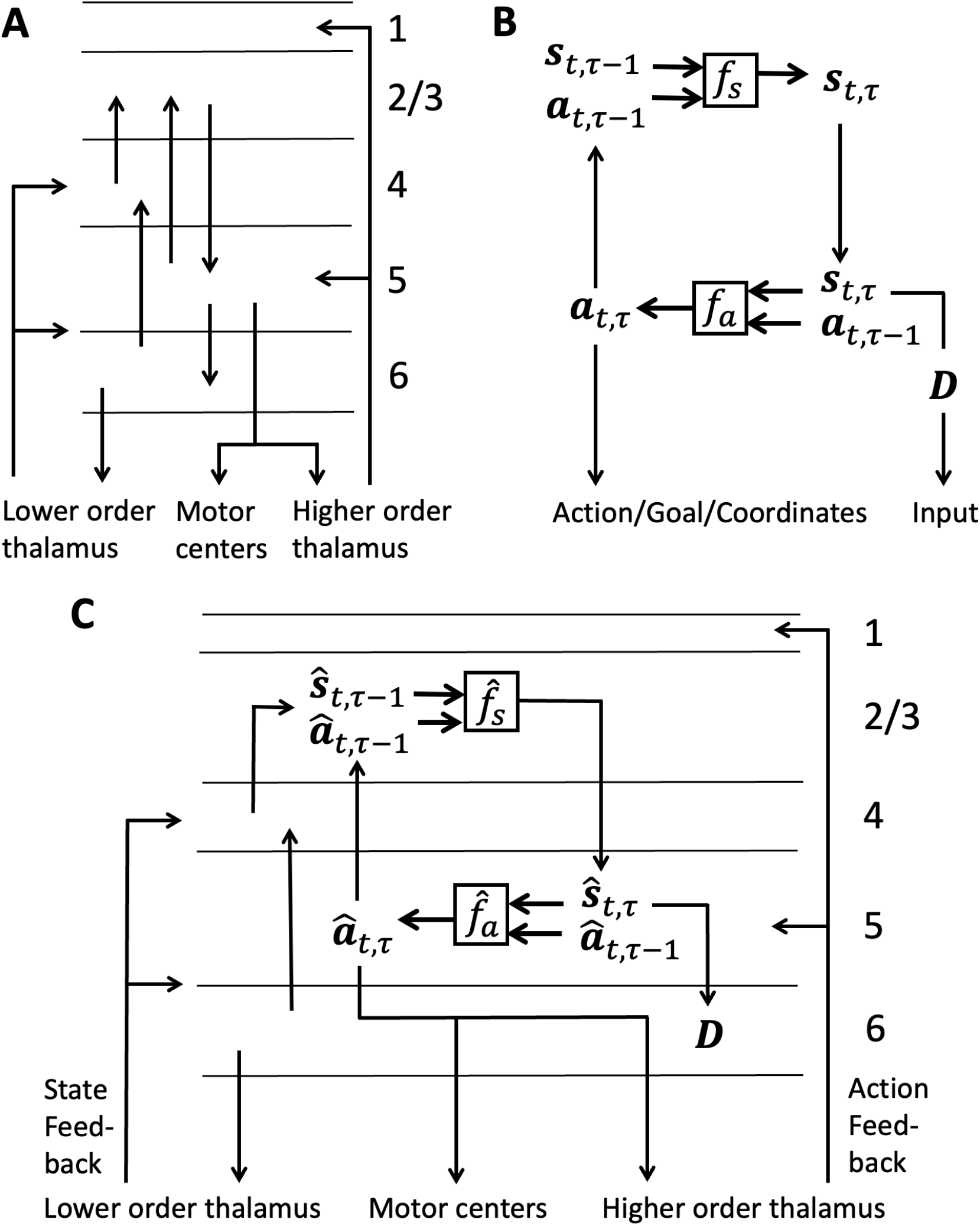
Canonical APC Cortical Module. (A) Depiction of the six-layered laminar structure of a cortical column, showing some of the major connections between layers and with the thalamus (not all connections are shown). (Based on [4, 18, 19]). (B) Canonical APC generative model. (C) One possible implementation of inference in a canonical APC cortical module for the generative model in (B). See text for details.

### 2.2 Computational Motivation

Normative computational considerations point to maintaining a close link between actions and their sensory consequences. For example, in model-based reinforcement learning [20] and more generally, in the framework of partially observable Markov decision processes (POMDPs) [21, 22], an intelligent “agent” interacts with the world by executing an action *a*_*t*_ at time *t* and this causes the agent’s “state” to change from *s*_*t*_ to *s*_*t*+1_; this change is governed by the state transition function *f*_*s*_(*s*_*t*_, *a*_*t*_) which generates an *s*_*t*+1_ according to a probability distribution *P* (*s*_*t*+1_ | *s*_*t*_, *a*_*t*_). The concept of state here captures the very general notion of hidden aspects of the world important to the agent, e.g., the location of the agent in a building to be navigated or a part of an object to be recognized such as the wheel of a vehicle. The agent does not have direct access to the state but the state is “partially observable” in that the agent has sensors which provide an “observation” about the state, e.g., a visual image of the room the agent is in or a (potentially obstructed) view of the wheel of the vehicle. When the agent executes an action, e.g., moving its body by walking or making an eye movement, the hidden state changes to the next state (a new location in the building or a new part of the vehicle such as the door). Estimating these hidden states as the agent makes movements is the essence of perception: both Bayesian inference models [23, 24] and more recent deep-learning models [25] attempt to solve this state estimation problem (albeit typically without movements in the case of static object recognition).

### 2.3 Canonical APC Cortical Module

#### 2.3.1 State Transition Function

The state transition function *f*_*s*_ described above is determined by the physics of the environment and the agent. By learning an approximation 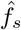 to *f*_*s*_ from its interactions with the world, the agent can learn an “internal model” of the world which can be used to run simulations of the world, imagine novel scenarios and explore what happens when particular actions are executed, and plan actions that lead to desirable states. The internal model 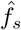 can be learned by randomly selecting actions and training the model on the resulting sensory data but a more realistic and less risky approach may involve guided exploration based on prior knowledge, imitation, instruction or strategies for balancing novelty-seeking versus risks and benefits. Although a Markovian single-level 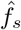 is too simplistic an internal model for most real-world problems, it can serve as a useful building block for more sophisticated hierarchical internal models as described below.

#### 2.3.2 Policy Function

An internal model (such as 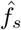 above) can be used to plan actions by unrolling the model into the future to explore the consequences of various action choices, but this mode of selecting actions requires considerable effort and deliberation (cf. type 2 or “system 2” thinking [26]). A much more efficient way to select actions (cf. type 1 or “system 1” thinking [26]) is to have a “policy” [20] 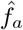, which maps the current state directly to an action that enables the agent to achieve the current goal. A policy 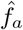 can be learned in several ways, e.g., by trial-and-error (“reinforcement learning” in the AI literature [20]) or more efficiently, via some combination of trial-and-error with instruction, imitation or planning.

#### 2.3.3 Coupling Perception and Action

Given a policy 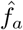 for a particular task or goal, the agent can execute actions from the policy while simultaneously predicting the sensory consequences of each action by sending an “efference copy” or “corollary discharge” to the learned model 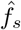. The agent can then correct its prediction of the new state of the world based on the new sensory observation that resulted from taking the action. The corrected estimate of the state *ŝ* can in turn be fed as input to the policy 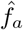 to generate the next action *â* for the task, continuing until the goal is achieved or the task times out.

#### 2.3.4 Generality and Flexibility

The generality of the above architecture can be appreciated by recognizing that such a sensory-motor generative model can generate (i) a sequence of images, mimicking the foveal images seen by an animal making eye, head or body movements, (ii) a sequence of touch sensations, modeling the process of using using hand movements to feel the shape of an object in the dark, (c) a sequence of odor sensations while navigating, (d) a sequence of proprioceptive sensations while walking, and (e) a sequence of abstract symbols, such as a “parent” or a “daughter”, while navigating a family tree (using abstract “actions” to go up or down the tree).

#### 2.3.5 Reference Frames

The sensory-motor generative model discussed above defines a *reference frame* [7, 27, *28] – it specifies how a particular object or abstract concept can be defined as a set of components that are related to each other in a specific way, e*.*g*., *relative location, orientation, or semantic relationship. For example, a physical object such as a car can be described as a set of parts (such as wheels, hood, doors etc*.*) at different locations and transformations within an object-centered reference frame. This can be captured by a state transition function that generates parts and a policy function that generates locations/transformations for the parts. In this case, the “actions” can be interpreted as coordinates within a reference frame*. A building or house to be navigated can similarly be described in terms of the different rooms and their connectivity, which once again can be captured by a transition function coupled with a policy for visiting the rooms. Finally, an abstract concept such as a family tree can be described in terms of a set of people and how they are connected via parent-child relationships, which again can be expressed by a transition function capturing these relationships (in terms of an abstract state and action generating another abstract state), and a policy for traversing and generating such a tree.

### 2.4 Potential Neuroanatomical Implementation

The above computational considerations and the generality of the framework motivate our canonical APC generative model shown in Figure 1B. The corresponding model for inference and learning, referred to in this paper as a *canonical APC cortical module*, is shown in Figure 1C, which also suggests one possible functional mapping of APC’s computational elements onto the cortical laminar structure in Figure 1A.

As shown in Figure 1C, the superficial layer neurons in cortical column, which receive the filtered sensory inputs from layer 4 and are recurrently connected to each other, are well-suited to implementing the statetransition function 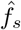. The class of motor output layer 5 neurons, which are also recurrently connected to each other, appear to fit the role of neurons computing the action/policy function 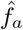. The other class of Layer 5 neurons, which convey information to other cortical areas and the striatum, could maintain the current state estimate *ŝ*_*t*_ by integrating the state prediction from layer 2/3 and correcting it with prediction errors from the thalamus. Layer 6 neurons receiving inputs from these state-estimating Layer 5 neurons are well-placed to compute the prediction for a lower-level area: at the lowest level, Layer 6 neurons predict sensory input *Īt* for the input *I*_*t*_ (Layer 6 neurons at a higher level would predict the cortical state at a lower level – see Section 3).

The thalamus is an integral part of cortico-cortical and cortico-subcortical computation [4]. We suggest that each feedforward pathway from the thalamus to Layers 4 and 5/6 of a cortical area conveys prediction errors. This was first suggested in the original Rao-Ballard predictive coding model for the LGN to Layer 4 primary visual cortex pathway [1]. We generalize this suggestion to include second-order thalamic nuclei such as the pulvinar. In this case, we suggest the Layer 5 inputs to the thalamus from a lower cortical area convey the current state estimate *ŝ*_*t*_ while the Layer 6 feedback from a higher cortical area to the same thalamic region conveys the prediction 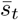; the interactions of these cortical estimates and predictions with inhibitory interneurons in the TRN and thalamus may enable computation of prediction errors, which are conveyed back to the cortex for updating cortical estimates of states and actions.

Additionally, the Layer 5 motor output neurons, e.g., those sending outputs to the superior colliculus, also send axon collaterals to higher-order thalamic nuclei. The same nuclei also receive motor information from subcortical motor centers such as the superior colliculus regarding actions executed. These thalamic nuclei are therefore in an ideal position to compare the actual action *a*_*t*_ (from superior colliculus or other motor center) and the cortical prediction *â*_*t*_. The resulting action feedback (e.g., in the form of action prediction errors) can be conveyed by the thalamus back to the cortex to enable the state transition network 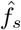 to correct its state prediction based on the actual action executed. Indeed, it is known that higher-order nuclei such as the pulvinar receive cortical layer 5 inputs and information from the superior colliculus, and send axons to superficial layers of area V1: such a circuit can explain, for example, the response of V1 neurons to saccadic eye movements [29].

The proposed neural implementation of APC above could potentially explain other neurophysiological results as well. For example, the prediction of next coordinates of a stimulus as a function of current action (via 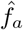) could explain how cortical circuits are able to anticipate the retinal consequences of eye movements and predictively update the retinal coordinates of stimuli [30]. The APC model’s neural implementation of reference frames may offer an explanation for object-centered parts-based representations in cortical areas such as V4 [31]. The laminar mapping suggested above is also consistent with previous predictive coding results suggesting prediction error-like responses in superficial layers and state-estimation-like responses in deeper layers [32]. The idea of Layer 5 output neurons encoding goals of movements is also consistent with the more specific proposal that M1 layer 5 neurons specify goals for movements rather than motor commands [33]. We discuss other implications and open questions pertaining to the neural implementation of the APC model in the Discussion section (Section 6).

## 3 Hierarchical Active Predictive Coding

### 3.1 Computational Motivation

Consider the problem of going to the grocery store from your house to buy some milk (Figure 2A). At a high level of abstraction, you may divide the task into sub-tasks (or “sub-goals”) such as walking to the door of your house from whichever room you are currently in, opening the door and walking to your garage where your car is parked, getting into car and driving to the grocery store, parking your car, walking to the entrance of grocery store, etc. Note that many of these sub-tasks can be *re-used* for solving a range of other problems (e.g., going to work, going to visit a friend, going to a restaurant etc.) and therefore, it may be useful to learn and maintain a policy for the sub-task (see previous section) to avoid planning actions each time the sub-task is re-used as part of a different task.

**Figure 2:**
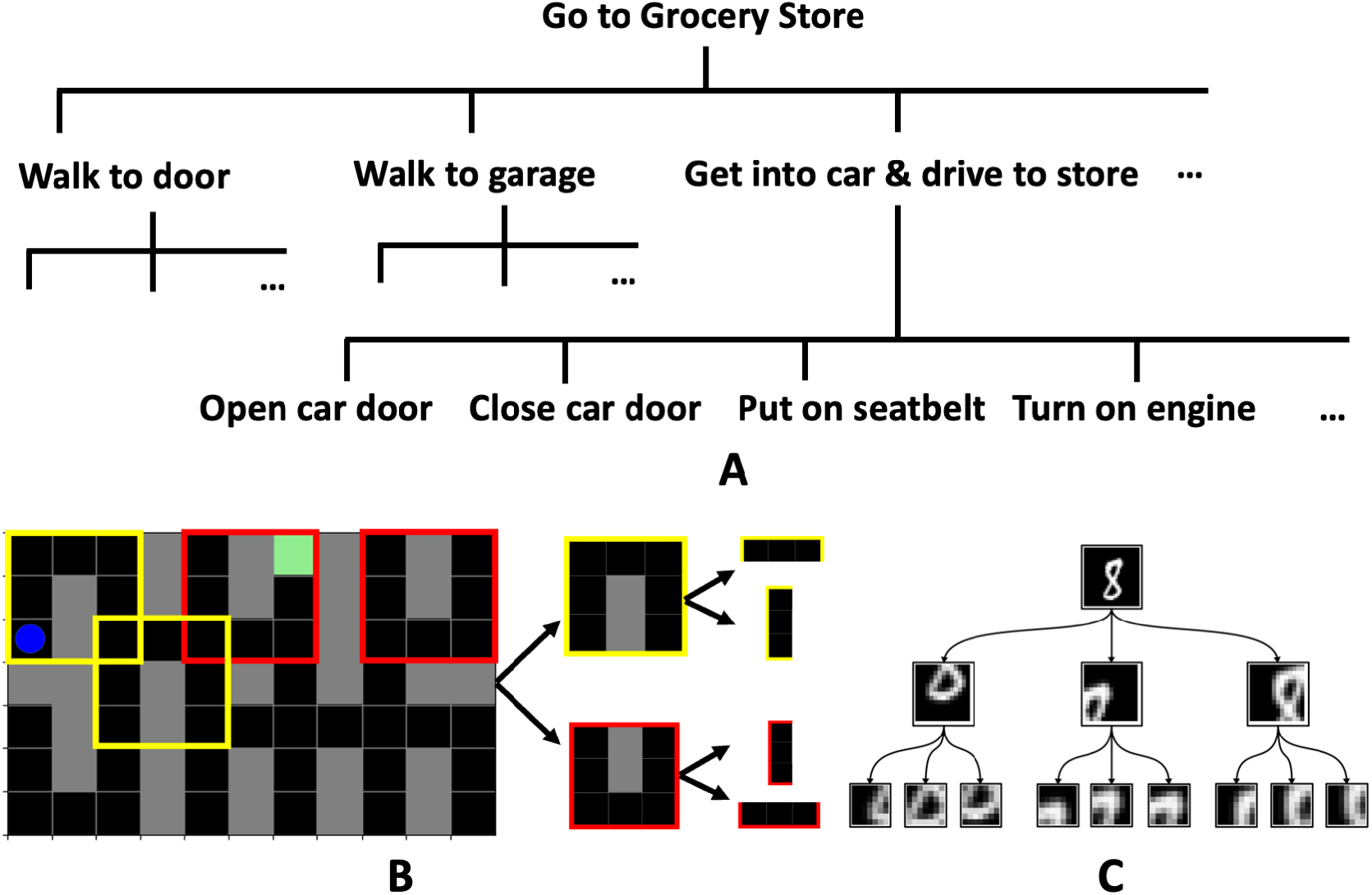
Modeling Complex Worlds and Tasks by Exploiting Compositionality. (A) Decomposition of the “Go to Grocery Store” problem into sub-goals/tasks, each of which can be further divided into sub-sub-goals/tasks. Note that the rate of change is faster at the lower levels compared to the higher levels, leading naturally to a temporal hierarchy. (B) A navigation problem in a maze-like building environment with corridors (black) and walls (gray), the blue dot indicating current location and green square the desired goal location. The structure of the environment can be understood in terms of its state transition dynamics, which in turn can be divided into the simpler transition dynamics of its compositional elements, two rooms outlined in yellow and red that appear at several different locations within the reference frame of the environment. These simpler elements can be further decomposed into horizontal and vertical corridors shown on the right that appear at different locations within the local reference frame of each room. (C) An object (such as a handwritten digit “8”) can be divided into parts (loops and curves at the middle level), each of which can be divided into sub-parts (strokes, lines, smaller curves at the lower level). Each part/sub-part is associated with its coordinates (location/transformation) within a local reference frame.

#### 3.1.1 Compositionality

The ability to divide a problem into components and re-use the components to solve new problems gets to the heart of *compositionality*, a powerful attribute of representations that may form the basis for the flexibility and fast generalization ability of human cognition [34, 35]. The APC model is motivated by the goal of learning compositional representations in a cortically-inspired neural architecture.

Consider our example task above of going to the grocery store. We can go beyond the division of this task into sub-tasks: each sub-task can in turn be divided into “sub-sub-tasks” operating at a lower level of abstraction. For example, as shown in Figure 2A, the sub-task of getting into your car and driving to the grocery store involves opening the door of the car, closing the door, putting on the seatbelt, turning on the engine, putting your foot on the brake pedal, shifting the car into “Drive” mode etc. Each of these sub-sub-tasks can in turn be decomposed into smaller tasks at an even lower level of abstraction, e.g., opening the door of the car may involve reaching towards the door handle, grasping the handle, pulling the handle, etc. The lowest level of abstraction, where the state changes happen at the fastest time scale, may involve activating muscles of the arm and hand in appropriate ways to enable the desired reaching and grasping for opening the door.

This strategy of tackling complex problems by division into sub-tasks, sub-sub-tasks and so on involves not only dividing a complex action into simpler actions but also dividing a complex state, defined by its state transition function, into simpler state transition functions. This is illustrated in the example in Figure 2B which is a simplification of the “Go to grocery store” problem: here the problem is to navigate in a relatively complex maze-like building environment with corridors (black) and walls (gray) (blue indicates the current location and green the current goal location). This environment can be divided into its compositional elements, two simpler rooms outlined in yellow and red that appear at different locations as shown in Figure 2B. These simpler rooms can in turn be decomposed into horizontal and vertical corridors as shown in the figure. Note that the rooms are “simpler” in the sense that they have simpler state transition functions than the original building environment, and the corridors in turn have simpler state transition functions than the rooms.

#### 3.1.2 Nested Reference Frames

Interestingly, the same concept of decomposing a task into sub-tasks, sub-sub-tasks, etc. as described above can be applied to the visual perception problem of representing an object in terms of parts, subparts etc. For example, a person can be represented in terms of the parts: head, arms, torso, and legs; a leg in turn is made up of sub-parts such as the foot, ankle, lower leg, knee, upper leg etc. Similarly, a handwritten digit, e.g., an “8”, can be divided into parts (loops and curves), each of which can be divided into sub-parts (strokes, lines, smaller curves) as shown in Figure 2C.

While an object can be defined in terms how its parts are located relative to each other (coordinates of the parts) within the object reference frame, each part can be similarly defined in terms of where the sub-parts are located within *that part’s reference frame*. This motivates the use of *nested reference frames*, where the reference frame at each abstraction level can be defined and used as a modular unit independent of how other reference frames are defined. This bestows on the nested representation substantial combinatorial flexibility in representing and solving complex tasks and problems in terms of simpler, re-usable components. The fact that the world we live in and the problems we seek to solve are amenable to compositional solutions and representations makes such an approach attractive, both from a computational and evolutionary perspective, and motivates the APC model’s hierarchical architecture.

### 3.2 Hierarchy

Nested reference frames and hierarchical states and actions are implemented in the APC model as follows (Figure 3A): a state representation vector *s*^(*i*+1)^ at abstraction level *i* + 1 generates a state transition *function* 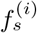 at the lower level *i* (along with an initial state vector 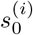 to start the lower-level state-action sequence); similarly, a higher-level action vector *a*^(*i*+1)^ generates a lower-level policy *function* 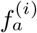 (called an “option” in reinforcement learning [36]). The two lower-level functions interact with each other, as in the canonical APC module (Figure 1B), to generate lower-level states and actions as shown in Figure 3A. Each such state and action can in turn generate transition and policy functions at an even lower level of abstraction. A lower-level sequence executes for a period of time until a condition is met (e.g., a sub-goal is reached, a task is completed or times out, or there is an irreconcilable error at that level). Then, control returns to the higher-level, which transitions to a new higher-level state and a new higher-level action, and the process continues. It is clear that such a generative model can generate the dynamics of the states (the “physics” of the world) and action sequences (produced by a policy) at different time scales, providing a mathematical and hierarchical way of describing complex tasks such as going to the grocery store.

**Figure 3:**
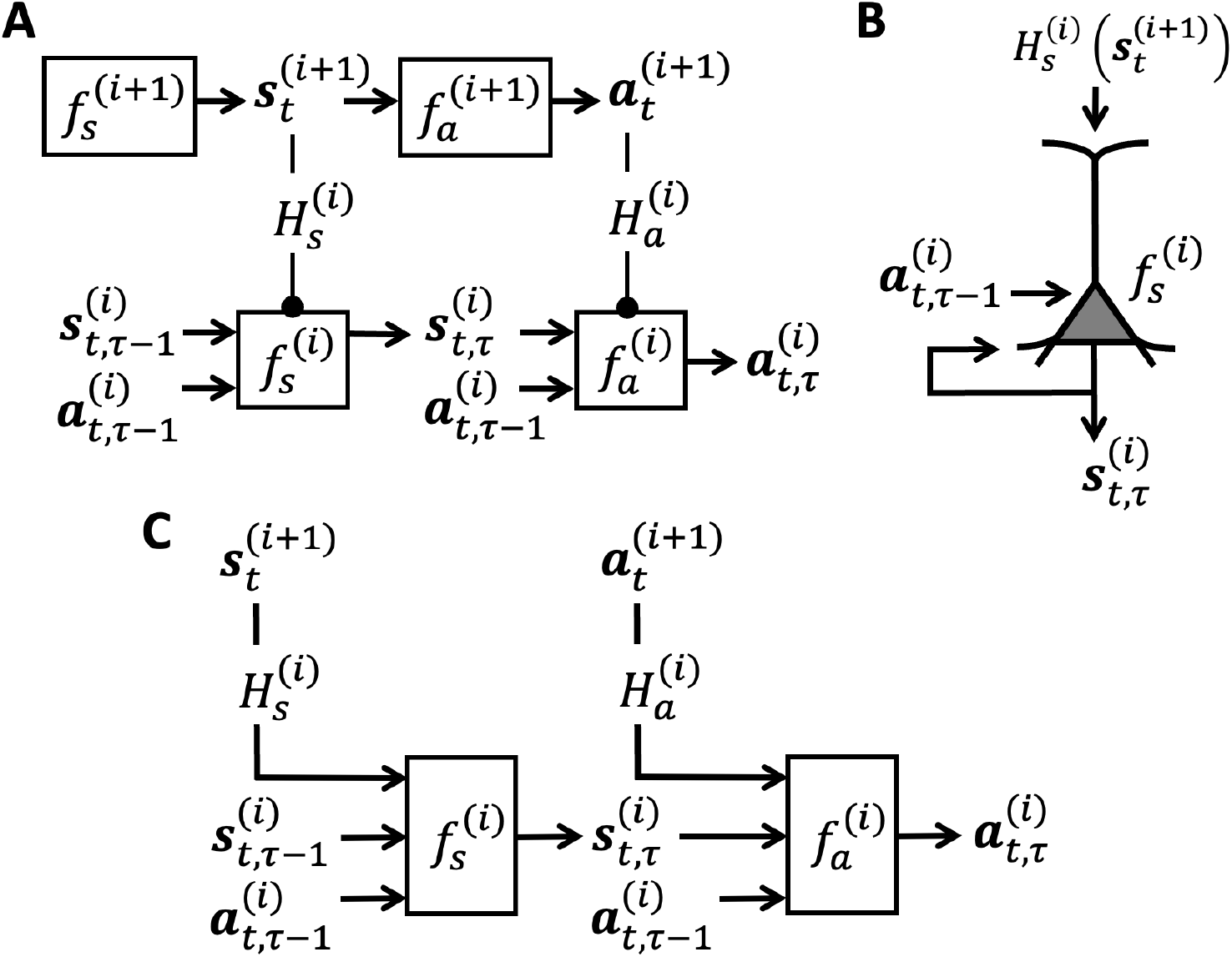
Hierarchical APC Model. (A) The diagram shows two levels (level *i* + 1 and *i*) of a hierarchical APC generative model. Each level has a state transition function *f*_*s*_ capturing the dynamics of the world at a particular level of abstraction, and a policy function *f*_*a*_ specifying that level’s actions/goals/coordinates (conditioned on the current highest level goal). The higher level state and action vectors at time *t* generate, via top-down networks *H*_*s*_ and *H*_*a*_, state transition and policy functions for the lower level, allowing the higher level to compose a sequence of states and actions at the lower level to accomplish a goal or sub-goal. (B) As depicted for a single pyramidal neuron here, we hypothesize that top-down inputs 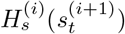 from a higher cortical area to the apical dendrites of a pyramidal neuron in a lower area modulate the dynamics of a network of such neurons (cf. gain modulation [37, 38]), allowing the higher area to change the function *f*_*s*_ or *f*_*a*_ being computed by these neurons. (C) Embedding-input-based implementation of the model in (A) in which higher-level state and action neurons maintain estimates of state and action vectors (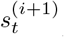 and 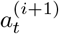) and modulate the lower-level state and action networks via top-down embedding inputs 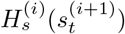 and 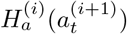 respectively.

#### 3.2.1 Neural Challenge and Possible Implementation

How can the hierarchical model in Figure 3A be implemented in networks of neurons? More specifically, how can a population of neurons, representing for example the higher-level state vector *s*^(*i*+1)^, *generate a whole function* 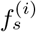 at the lower level?

One possible solution, from the field of AI, is to use a “hypernetwork” [39], which is a neural network which produces the synaptic weights of another neural network *P* (called the “primary network”). In our case, the higher-level state vector *s*^(*i*+1)^ would be fed as input to a network *H*^(*i*)^ which produces the weights for a lower-level network computing 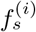. From a biological point of view, one may be tempted to dismiss such a model as biologically implausible since a biological neural network cannot create another neural network *ex nihilo*. However, instead of generating weights, a hypernetwork can also *modulate* the existing weights of the “primary” neural network *P*, e.g., by multiplying the synaptic inputs to *P* (or *P* ‘s outputs) by a gain factor, and by doing so, it can change the function being computed by the neural network *P*. Such “gain modulation” appears to be a ubiquitous phenomenon in the cortex and has been the subject of extensive investigations, both computationally and experimentally [37, 38, 40–44].

A second related solution, again suggested by AI research, is to use the “embedding approach” [45, 46]: the higher-level state vector *s*^(*i*+1)^ is fed as input to a network *H*^(*i*)^ that produces an *embedding vector e*. The embedding vector *e* is provided as an *additional non-changing contextual input* to a lower-level neural network *P* that computes 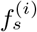. The higher-level can therefore control the function being computed at the lower level by changing the embedding vector input *e*. Such an approach has recently been used in a computational neuroscience study to show how a single recurrent network can be used to solve multiple tasks by changing its task input (or “rule input”) [46, 47]. This is a special case of the model we are suggesting; in our model, the embedding input can be a rich adaptive top-down input provided by a higher-level network, and this can be repeated to learn a hierarchy. Note that from previous AI research, the embedding approach and the hypernetwork approach can be shown to be equivalent in terms of computational function but the hypernetwork approach may offer greater efficiency under certain assumptions [45].

### 3.3 The Hierarchical APC Model

The computational and neural considerations above suggest the following hierarchical APC model. At each hierarchical level, the higher-level state and action vectors, *s*^(*i*+1)^ and *a*^(*i*+1)^ respectively, use gain modulation and/or embedding inputs to modulate the lower-level networks computing the lower-level state and action functions 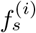 and 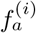 respectively, Such modulation of lower-level networks is consistent with the previously suggested modulatory role for layer 1 and superficial layer inputs to apical dendrites of cortical neurons (Figure 3B) [37].

### 3.3.1 Modulation of State Networks by Feedback to Model Complex Environmental Dynamics and Physics

Figure 3C shows an implementation of a 2-level APC network via the embedding approach (see [10, 12, 48] for hypernet-based implementations). As shown in Figure 3C (left), the higher-level (level *i* + 1) state neurons maintain an estimate of the state vector 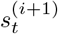 at time *t* and modulate the lower-level state network via top-down feedback given by the embedding input 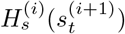.

The lower-level state neurons (at level *i*) maintain an estimate of the lower-level state 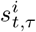 where *τ* denotes a time step at the lower level within the higher level time interval given by *t*. In the two-level network, this lower-level state makes a prediction of the input via a “decoder” network *D*. In the simplest case where *D* is a linear matrix *U*, this lowest level of APC is equivalent to the generative model using in sparse coding (*I* = *Us* where *s* is sparse [49]). More generally, *D* can be a 1-layer RELU network [48] or a multi-layer decoder [11, 12].

The state networks at all levels can be trained using *self-supervised learning* based on *prediction errors*: at each time step, the network predicts the next input as a function of previous state and action, and the prediction error can be backpropagated [10, 48] or used in more neurally plausible ways, e.g., using local errors [1], to update the weights of the state networks at all levels.

In addition to being used for self-supervised learning, prediction errors can also be used for inference to correct the estimates of state at each level using gradient-based updates similar to the original Rao-Ballard predictive coding model [1]: here feedforward pathways convey prediction errors to update state estimates and feedback pathways convey top-down modulation as described above (see [48] for an example based on modulatory hypernetworks).

Such an arrangement allows the network to learn internal models of complex environmental physics and dynamics using a “divide-and-conquer” compositional strategy: any complex state-transition function is modeled by the network as a composition of simpler state-transition functions operating over short timescales by modulating lower-level networks using top-down feedback. For example, the physical state changes that a hand goes through when opening a car door can be modeled as a sequence of brief muscle activation patterns. The car door itself goes through a state change from being closed to open, which can again be modeled as a sequence of simpler lower-level state transitions: the door handle transitions from one position to another, the angle of the door transitions from an angle of zero degrees to increasing larger degrees up to a maximum, etc. These states and state transitions can in fact be regarded as a *sensorymotor description of an object*, and can be used for object recognition. We provide illustrative examples of this idea in Section 4, where we apply it to recognizing visual objects based on sequences of their parts and sub-parts, and to navigating a complex environment by dividing it into smaller components.

#### 3.3.2 Modulation of Action Networks by Feedback to Solve Complex Problems via Compositionality

Figure 3C (right) shows the higher- and lower-level action networks that are coupled to their respective state networks. The higher-level (level *i* + 1) action neurons represent an abstract action (such as “open the door”) via an action vector 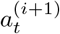 at time *t*. Note that the abstract action vector can also be interpreted as a *higher-level goal* (“achieve the state where the door is open”). Given the abstract action (goal) 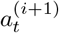 top-down feedback given by the embedding input 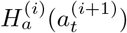 modulates the lower-level action network as described above, implementing the fine-grained details of the abstract action.

Specifically, as depicted in Figure 3C (right), the lower-level action network produces a lower-level action 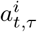 where *τ* denotes a time step at the lower level within the higher level time interval given by *t*. This action is fed back to the state network at the same level to generate the next state prediction, which generates the next action 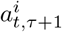 and so on until either the sub-goal for that level is achieved or the level times out and returns control back to the higher level.

The action networks at all levels are trained to compute policies (state-to-action mappings) to achieve the current overarching goal or task. There are multiple ways to train the action networks:

- **Planning**: The APC model incorporates state transition networks at all levels, providing it with a multi-scale internal model of the world. Therefore, the model can plan a sequence of actions to achieve a goal, starting from the highest abstraction level, and searching for sequences of actions that are likely to achieve the goal or move the state closer to the goal as predicted by the learned state transition functions. This strategy is consistent with active inference [50], planning by inference [51–53] and model predictive control [54] (which we illustrate in Section 4 in our example on planning for navigation). If a planned sequence of actions is successful in achieving a desired goal, these actions can be used as “labels” in supervised learning to train the policy or action network 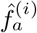 at all levels.
- **Reinforcement Learning**: An alternate way to train the action networks is to use hierarchical reinforcement learning [55, 56] in which each level explores sequences of actions and trains its action network to maximize the total expected reward it receives according to a reward function which may be specific to that level. For example, the higher level can be trained to generate a sequence of explicit sub-goals for the lower level and the lower level then receives (intrinsic) rewards according to how close the lower level state gets to each sub-goal set by the higher level [57]. We provide an illustrative example of a reinforcement learning-based method for training an APC model’s action networks in Section 4.
- **Policies providing Priors for Planning**: The APC model incorporates both a model of the world (the state transition networks 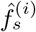 and policy networks 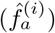 (conditioned on the current highest level goal) that predict actions for any given state at each abstraction level *i*. These predicted actions can be regarded as priors, in a Bayesian sense, to guide the search for successful actions in planning. When a task in unknown or new, the policy network is not yet trained and the predicted actions will have high uncertainty. In this case, the policy network cannot bias the planner in a useful way and the planning algorithm has to search over a much broader space of actions, requiring much effort and deliberation (cf. “system 2 thinking” [26]). When the task is frequently encountered and has been repeatedly solved successfully, the policy networks 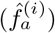 are well-trained and predict actions with high confidence and low uncertainty. As a result, the same planning algorithm can leverage the highly confident suggestions from the policy networks to restrict its action search space, needing little or no exploration to quickly select successful actions (cf. “system 1 thinking” [26]).^1^

In summary, the APC model’s hierarchical action networks 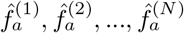 can be implemented by action neurons representing an abstract action 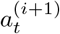 at a higher level modulating a lower-level action network 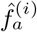. This top-down modulation changes the policy function this lower-level network is computing such that it produces a sequence of lower-level actions 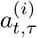 for achieving the higher-level goal (abstract action) represented by 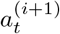 The action networks and the top-down modulation networks can be learned via planning, reinforcement learning or a combination of both.

A great advantage of learning a set of hierarchical actions as suggested above is compositionality: the abstract action vectors *a*^(*i*+1)^ (with its associated lower-level implementation via 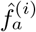) can be re-used and chained together by the planner with other abstract actions to solve new tasks. Going back to our example of going to the grocery store, the abstract action of walking to the door of your house, when successfully learned within a lower-level network 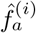, can be re-used for solving other problems such as going to work, going to a restaurant etc. We provide a concrete example of this compositional approach to problem solving used by the APC model in Section 4.

### 3.4 Fast Generalization via State and Action Embedding Spaces

We note that since both the abstract state vectors *s*^(*i*+1)^ and abstract action vectors *a*^(*i*+1)^ are continuous-valued vectors, there exists a potentially powerful opportunity to leverage the structure of these high-dimensional state and action embedding spaces. For example, by including smoothness constraints on these vectors during learning, it may be possible to use interpolation/extrapolation in this embedding space or sample in the neighborhood of known *s*^(*i*+1)^ and *a*^(*i*+1)^ to *generate on-the-fly* new state transition functions 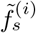 and new policy functions 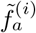, opening the door to fast generalization and transfer of knowledge to new problems. Some preliminary steps in this direction are explored in [12].

### 3.5 Potential Neuroanatomical Implementation

Building on the suggested implementation of the canonical APC cortical module in Figure 1C (Section 2.4), we propose that current state and action vectors *ŝ*^(*i*+1)^ and *â*^(*i*+1)^, which are estimated in a higher cortical area A, modulate the state and action networks 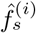 and 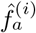 in a lower cortical area B via feedback connections that target the superficial and deep layers of area B [59]. It is known that feedforward connections that target Layer 4 of area A arise from area B and from the higher-order thalamic region receiving “driver input” from area B [4, 59]. We propose that these feedfoward connections carry the state and action feedback (e.g., prediction errors) that enable the higher area A to correct its abstract state and action estimates in case there is a significant error in the task, and generate new abstract state and action vectors for modulating the lower area B’s state and action networks to tackle the error.

## 4 Illustrative Examples: Diverse Functions from the Same APC Architecture

The APC architecture was inspired by the proposition that evolution may have replicated a common computational principle across cortex [5–7]. If that is the case, one would expect the same architecture to be able to solve problems as diverse as visual perception and planning movements in complex environments. In this section, we show that this is indeed the case by illustrating how the APC architecture can be used for both vision (recognizing visual objects from their parts, learning part-whole hierarchies) and planning (modeling navigation problems in terms of simpler navigation problems, exploiting hierarchical actions for efficient planning).

### 4.1 Example 1: Active Visual Perception

#### 4.1.1 Recognizing Objects and Learning their Parts using Eye Movements

Human vision can be viewed as an active sensory-motor process that samples a scene via eye movements to move the high-resolution fovea to appropriate locations and accumulate evidence for or against competing visual hypotheses [60]. The APC model is well-suited to the sensory-motor nature of human vision, given its integrated state and action networks. We briefly review here our recent simulation study of the APC model applied to the active vision and part-whole learning problem [10, 11].

We simulated a 2-level APC model, with top level and bottom level state and action vectors. The active nature of human vision as follows: actions in the APC model emulated eye movements (or “attention”) and corresponded to moving a “glimpse sensor” [61] which extracts high-resolution information about a small part of a larger input image. Given an input image *I* (of size *P* × *P* pixels), this sensor, *G*, takes in a location *l* and a fixed scale fraction *m*, and extracts a square glimpse/patch *g* = *G*(*I, l, m*) centered at *l* and of size (*mP*) × (*mP*). The location *l* to fixate next is selected by the bottom level action network within the reference frame specified by the top level. The bottom state network generates a prediction of the glimpse and the prediction error is used for updating the state and for learning.

In more detail, at each “macro-step” *t*, the top-level action vector *A*_*t*_ generates two values: (a) a location *L*_*t*_ and (b) a “macro-action” (or option) *z*_*t*_. The location *L*_*t*_ is used to restrict the bottom level to a sub-region 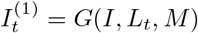 of scale *M* centered around *L*_*t*_, corresponding to a new frame of reference selected by the top level within the input image. The option *z*_*t*_, which operates over this frame of reference, is used as an input to the top-down network *H*_*a*_ to generate the embedding input to the bottom level policy/action RNN. For exploration during reinforcement learning of the action network, we treat the output of the location network as a mean value 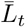 and add Gaussian noise with fixed variance to sample an actual location: 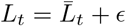, where *ϵ* ∼ N (0, *σ*^2^). We do the same for the option *z*_*t*_.

Based on the current bottom-level state and action, the bottom-level action RNN generates a new action *a*_*t,τ*_. A location *l*_*t,τ*_ is chosen as a function of *a*_*t,τ*_, resulting in a new glimpse image 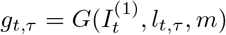 of scale *m* centered around *l*_*t,τ*_ and yielding a nested reference frame within the larger reference frame of 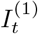 specified by the higher level. The bottom level follows the same Gaussian noise-based exploration strategy for sampling locations as the top level. The bottom-level state vector *s*_*t,τ*_, along with locations *L*_*t*_ and *l*_*t,τ*_, are fed to a generic decoder network *D* to generate the predicted glimpse *ĝ*_*t,τ*_. Following the predictive coding model [1], the resulting prediction error *ϵ*_*t,τ*_ = *g*_*t,τ*_ − *ĝ*_*t,τ*_ is used to update the state vector: *s*_*t,τ*+1_ = *f*_*s*_(*s*_*t,τ*_, *ϵ*_*t,τ*_, *l*_*t*_; *θ*_(*s*)_(*t*)). For the results below, the state networks at both levels were trained to minimize image prediction errors while the action networks were trained using reinforcement learning (the REINFORCE algorithm [62]).

We tested this APC network on the task of sequential part/location prediction and image reconstruction of objects in the following datasets: (a) MNIST: Original MNIST dataset of 10 classes of handwritten digits; (b) Fashion-MNIST: Instead of digits, the dataset consists of 10 classes of clothing items; (c) Omniglot: 1623 hand-written characters from 50 alphabets, with 20 samples per character. For this APC simulation study, we used 3 macro- and 3 micro-steps (except 4 macro-steps for Omniglot). A single dense layer, together with an initial random glimpse was used to initialize the state and action vectors of the top level (see [10] for details).

Figure 4 shows an example of a learned parsing strategy by the two-level APC model for an MNIST digit. The top-level learned to select actions that cover the input image sufficiently while the bottom level learned to parse sub-parts inside the reference frame computed by the higher level. Note how the initially uncertain hypothesis (blurry “0”) is refined as the model makes “eye movements” to accumulate evidence and converge on the identity of the input digit.

**Figure 4:**
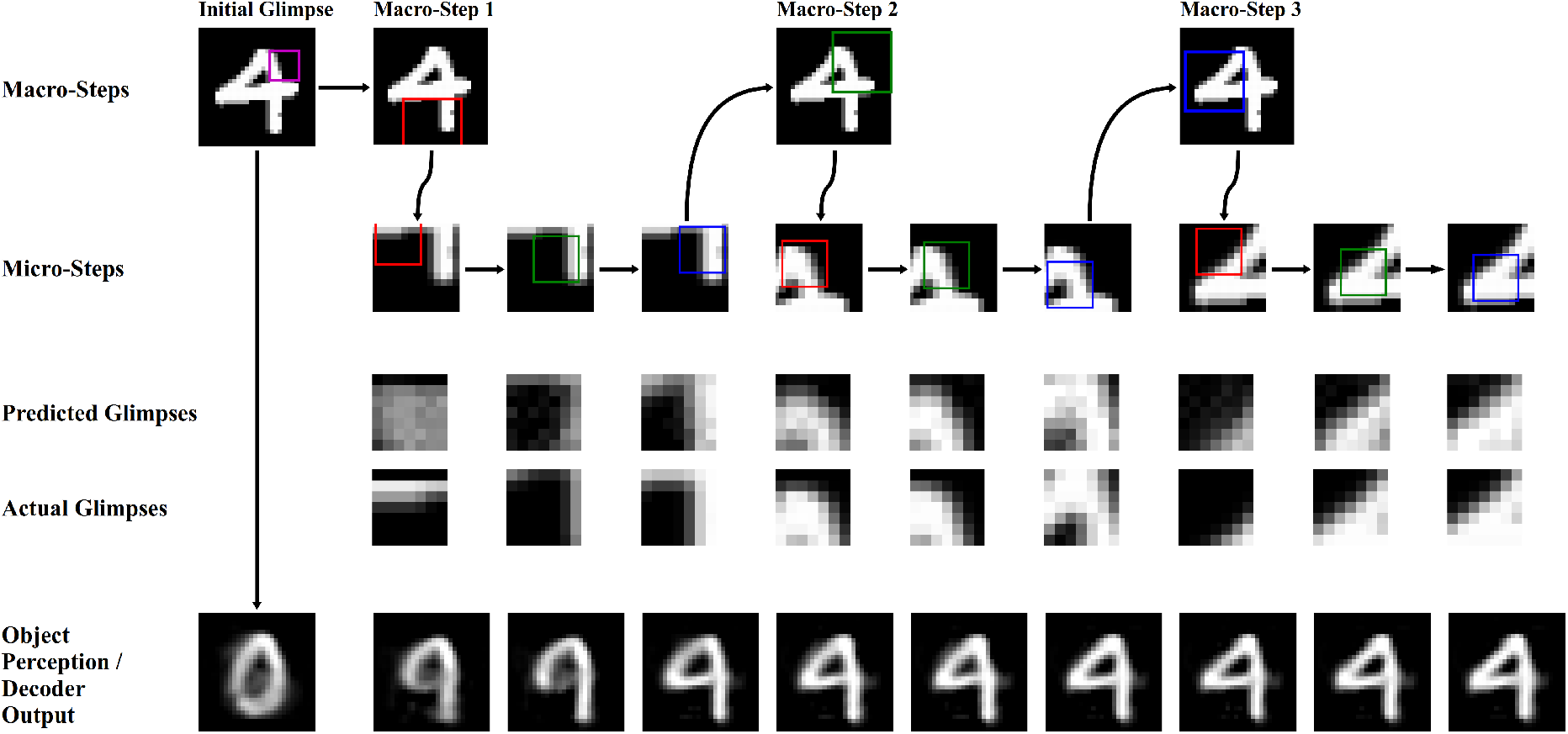
Hierarchical Parsing of an Image by the APC Model and an Illustration of Perceptual Stability in the presence of Eye Movements: 1st row: Initial glimpse (purple box) and higher-level reference frames selected by the model for this image (red, green, blue boxes), 2nd row: Lower level image regions visited by the model within each top-level frame, 3rd & 4th rows: Predicted versus actual parts/”glimpses” of the image, and 5th row: “Perception” of the model (object reconstructed by a decoder network from current network state) over time. Note the “perceptual” stability in the model despite jumps in the sampled glimpses (see row of Actual Glimpses) as the object hypothesis is gradually refined based on accumulated evidence.

#### 4.1.2 Perceptual Stability despite Eye Movements

Figure 4 also suggests a potential explanation for why human perception can appear stable despite dramatic changes in our retinal images as our eyes move to sample a scene: the last row of the figure shows how the model maintains a visual hypothesis that is gradually refined and which does not exhibit the kind of rapid changes seen in the sampled image regions (“Actual Glimpses” in Figure 4).

#### 4.1.3 Reference Frames and Part-Whole Hierarchy Learning

Figure 5 shows a learned part-whole hierarchy for an MNIST input, in the form of a parse tree of parts and sub-parts (strokes and mini-strokes) along with their locations within nested reference frames. The model learns different parsing strategies for different classes of objects, as seen in Figure 6 for different clothing items from the Fashion-MNIST dataset.

**Figure 5:**
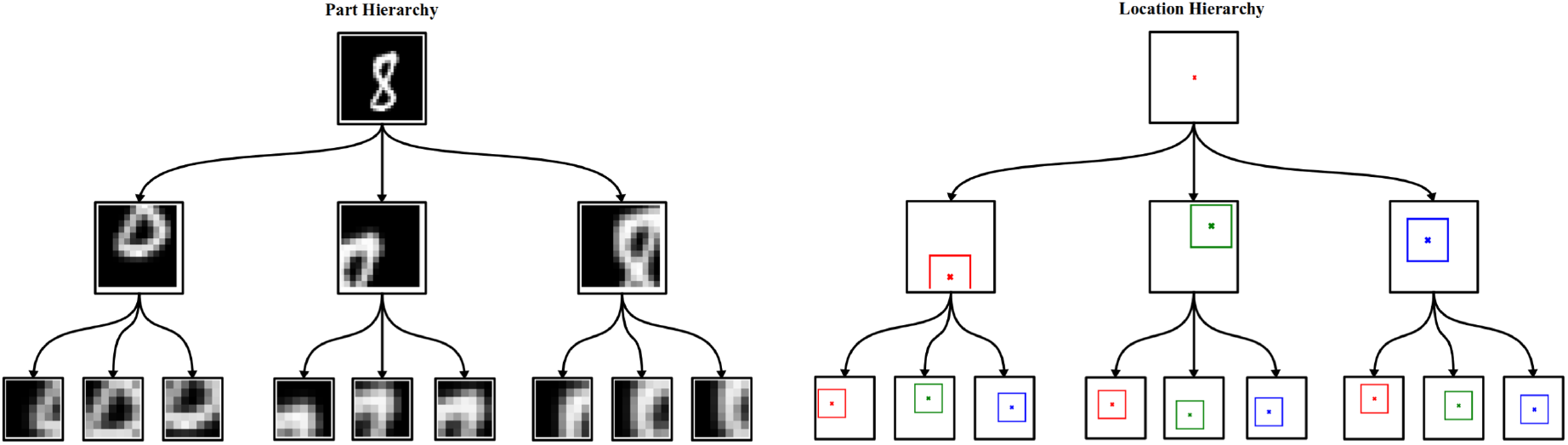
Compositional Part-Whole Hierarchy for an Object: The digit “8” is parsed using a parse tree of parts and sub-parts (left) and their corresponding coordinates (locations) within their respective reference frames (right). The representation is compositional because the same set of parts and sub-parts can be re-used at other locations and with other transformations to compose other digits that share these features.

**Figure 6:**
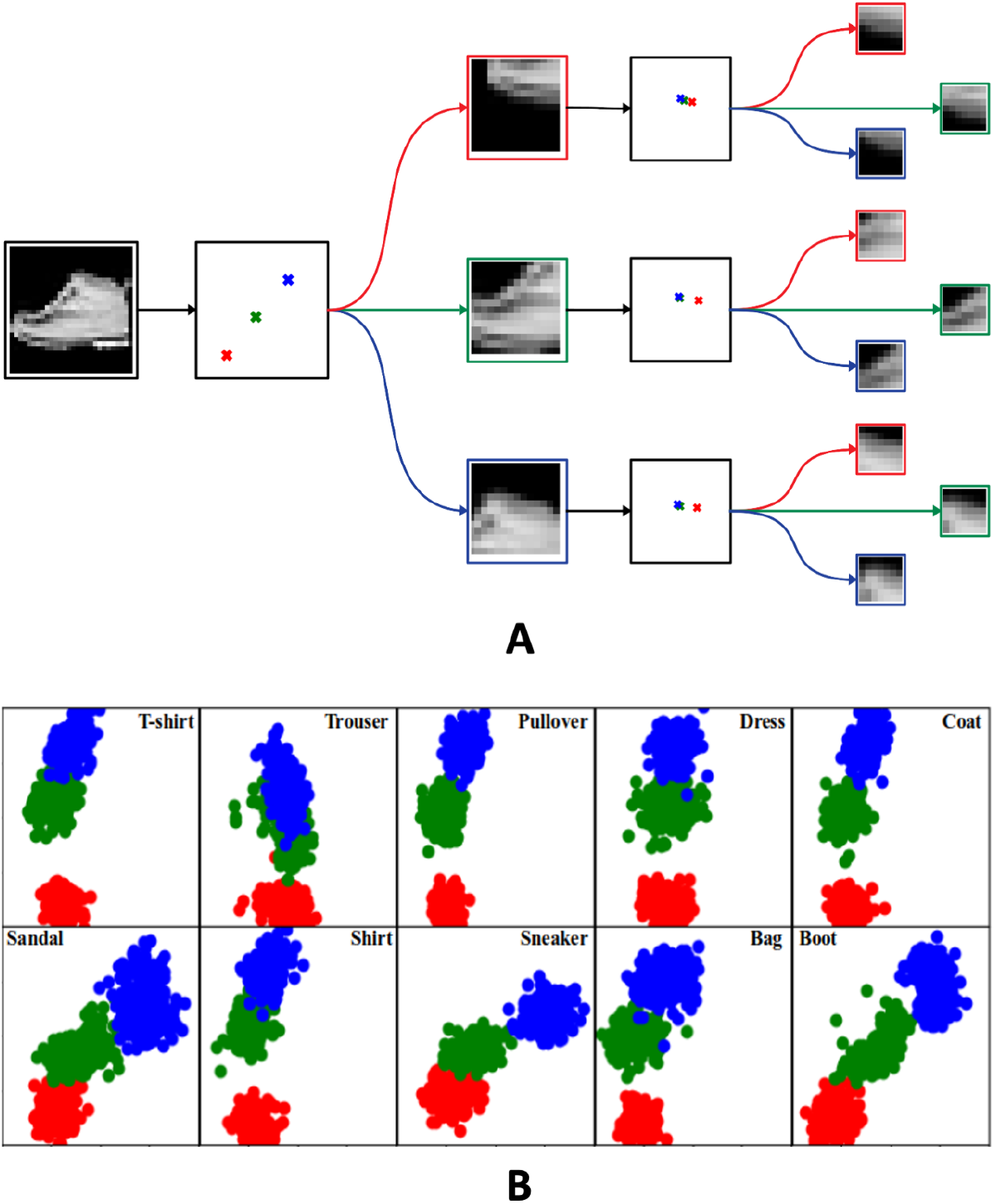
Class-Based Hierarchical Representation of Object Parts and Locations: (A) The diagram shows the parts and sub-parts recognized by a 2-level APC network trained on the Fashion-MNIST dataset for an input image of a sneaker. The red, green and blue dots show the average sampled locations for the “sneaker” class, visited in the following order within each frame of reference: 1st: red, 2nd: green and 3rd: blue. (B) Each panel shows the higher-level part locations selected by the trained APC network in (A) for all classes in the dataset (red, green, blue same as in A). Note the differences in the network’s action strategies between vertically symmetric items (shirts, trousers, bags) and footwear (sandals, sneakers, boots).

#### 4.1.4 Prediction of Parts, Pattern Completion and Transfer Learning

We investigated the predictive and generative ability of the trained model by having the model “hallucinate” different parts of an object. This was done by setting the prediction error input to the lower level network to zero, which disconnects the model from the input and forces it to predict the next sequence of parts to “complete” the object. Figure 7A shows that the model has learned to generate plausible predictions of parts given an initial glimpse.

**Figure 7:**
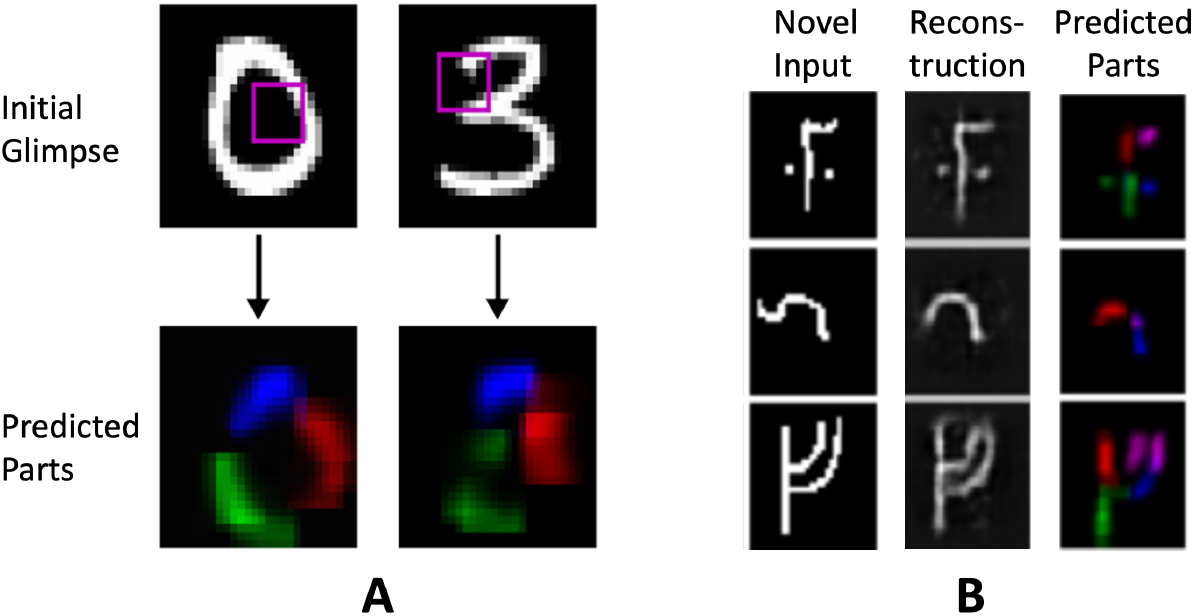
Prediction of Parts, Pattern Completion and Transfer Learning: (A) Given only an initial glimpse (purple box) for an input image (a “0” and a “3”), an APC model trained on the MNIST handwritten digits dataset predicts its best guess of the parts of the object and their locations (red, green and blue segments in row below). (B) APC model trained on the Omniglot handwritten characters dataset (from 50 different alphabets) can transfer its learned knowledge to predict parts of previously unseen character classes. First column: input image from a novel character class. Middle column: APC model’s reconstruction of the input. Last column: parts predicted by the model. (Adapted from [10]).

We also tested transfer learning in the APC model using the task of reconstruction of unseen character classes from the Omniglot dataset. We trained a 2-level APC model to reconstruct examples from 85% of classes from each Omniglot alphabets. The rest of the classes were used to test transfer: the trained model had to generate new state and action networks to predict parts for new character classes for each alphabet. The model successfully performed this transfer task (Figure 7B) (see [10] for quantitative results).

### 4.2 Example 2: Hierarchical Planning

#### 4.2.1 Modeling the “Physics” of Complex Environments in terms of Simpler Elements

We next briefly review our recent work [10] showing that the same APC framework used for learning part-whole hierarchies can also be used to to learn hierarchical world models that simplify the state-transition dynamics (“physics”) of a potentially complex environment into a sequence of simpler transition functions. Consider the problem of navigating from any starting location to any goal location in a large “multi-rooms” building environment such as the one in Figure 8A (gray: walls, blue circle: current location, green square: current goal location). In the traditional non-hierarchical reinforcement learning (RL) approach, the states are discrete locations in the grid, and actions are going north (N), east (E), south (S) or west (W). A large reward (+10) is received at the goal location, with a small negative reward (−0.1) for each action to encourage shortest paths.

**Figure 8:**
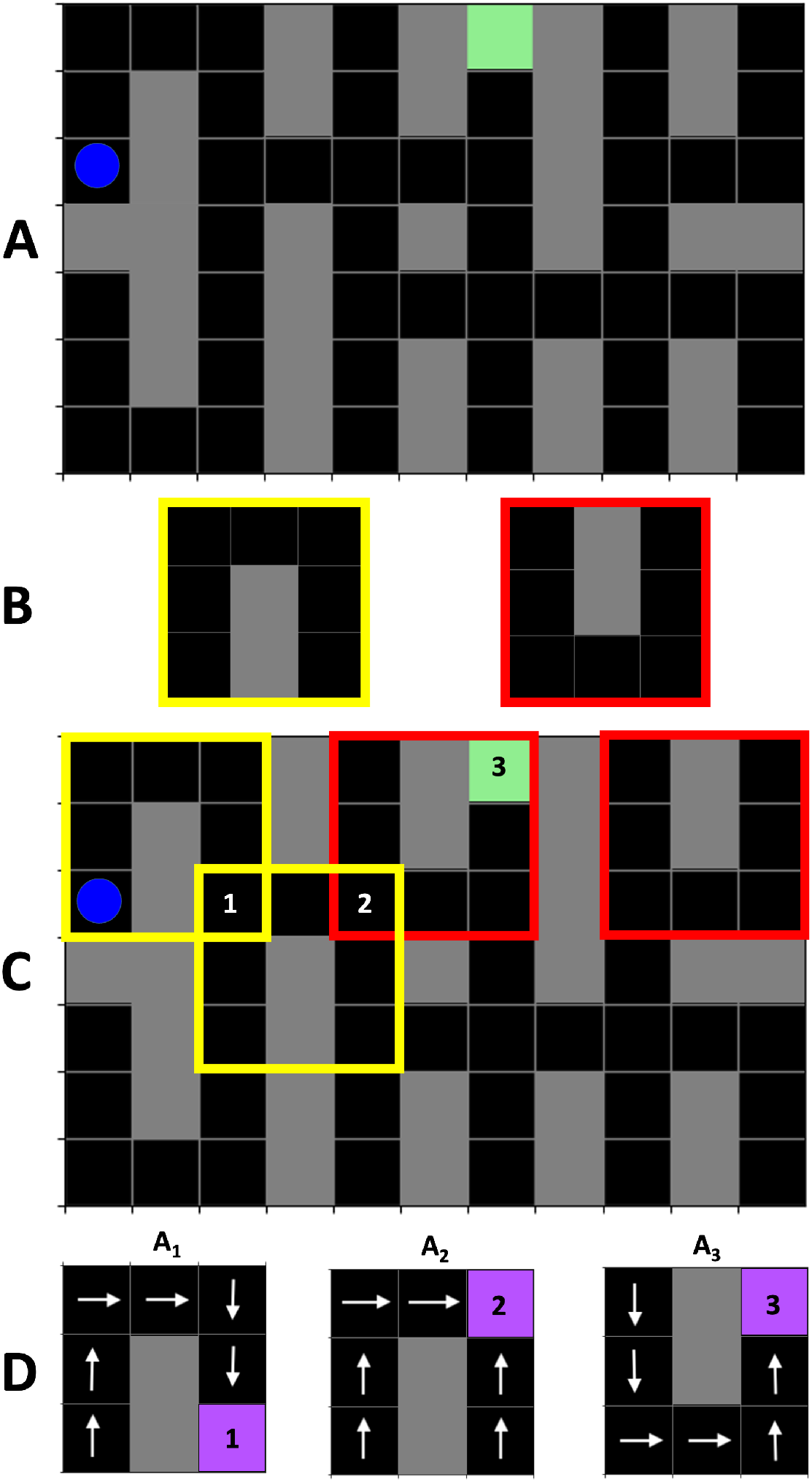
Solving a Navigation Task using a State-Action Hierarchy. (A) The problem of navigating in a large building can be reduced to planning using high-level states (B) and higher level actions ((C) and (D)). Blue: current location, gray: walls, green: current goal location. See text for details. (Adapted from [10]).

To simplify the transition dynamics of the larger environment, note that just as an object (e.g., an MNIST digit in Section 4.1.1) consists of the same parts (e.g., strokes, curves) occurring at different locations, the multi-rooms environment in Figure 8 is also made up of the same two components (“Room types” S1 and S2), shown in Figure 8B, occurring at several different locations (some example locations are highlighted by yellow and red boxes in Figure 8C). These components form part of the higher-level states in the APC and are defined by state embedding vectors S1 and S2, which can be trained to generate, via the top-down network *H*_*s*_ (Figure 3A), the lower-level transition functions 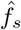 for rooms S1 and S2 respectively.

Similar to how the APC model was able to reconstruct an image in Section 4.1.1 using higher-level action embedding vectors to generate policies and actions (locations) to compose parts from sub-parts, the APC model can compute higher-level action embedding vectors **A**_*i*_ (option vectors) for the multi-rooms world that generate, via top-down network *H*_*a*_ (Figure 3A), lower-level policies 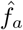 which produce primitive actions (N, E, S, W) to reach a goal *i* encoded by **A**_*i*_.

#### 4.2.2 Local Reference Frames allow Policy Re-Use and Hierarchical Planning

Figure 8D illustrates the bottom-level policies for three such action embedding vectors **A**_1_, **A**_2_ and **A**_3_, which generate policies for reaching goal locations 1, 2, and 3 respectively. Note that the **A**_*i*_ are defined with respect to higher-level state S1 or S2. Defining these policies to operate within the local reference frame of the higher-level state S1 or S2 (regardless of global location in the building) confers the APC model with enormous flexibility because the *same policy can be re-used at multiple locations* to solve local tasks (here, reach sub-goals within S1 or S2). For example, to solve the navigation problem in Figure 8C, the APC model only needs to plan and execute 3 higher-level actions or options: **A**_1_ followed by **A**_2_ followed by **A**_3_, compared to planning a sequence of 12 lower-level actions to reach the same goal.

The two-level APC model was trained as follows. The higher-level states represented 3 × 3 local reference frames and were defined by an embedding vector generating the transition function for “room type” S1 or S2, along with the location for this local reference frame in the global frame of the building. The lower-level action network was trained to map a higher-level action embedding vector **A**_*i*_ to a lower-level policy for navigating to a particular goal location *i* within S1 or S2. For this study, eight embedding vectors **A**_1_, …, **A**_8_ were trained, using REINFORCE-based RL [62], to generate via the network *H*_*a*_ eight lower-level policies to navigate to each of the four corners of room types S1 and S2.

The higher-level state network was trained to predict the next higher-level state (decoded as an image of room type S1 or S2, plus its location) given the current higher-level state and higher-level action. This trained higher-level state network was used for planning at each step a sequence of 4 higher-level actions using random-sampling shooting model predictive control (MPC) [54]: random state-action trajectories of length 4 were generated using the higher-level state network by starting from the current state and picking one of the four random actions *A*_*i*_ for each next state; the action sequence with the highest total reward was selected and its first action was executed, and this process was repeated. Figures 9A-C illustrate this high-level planning and MPC process using the trained APC model.

**Figure 9:**
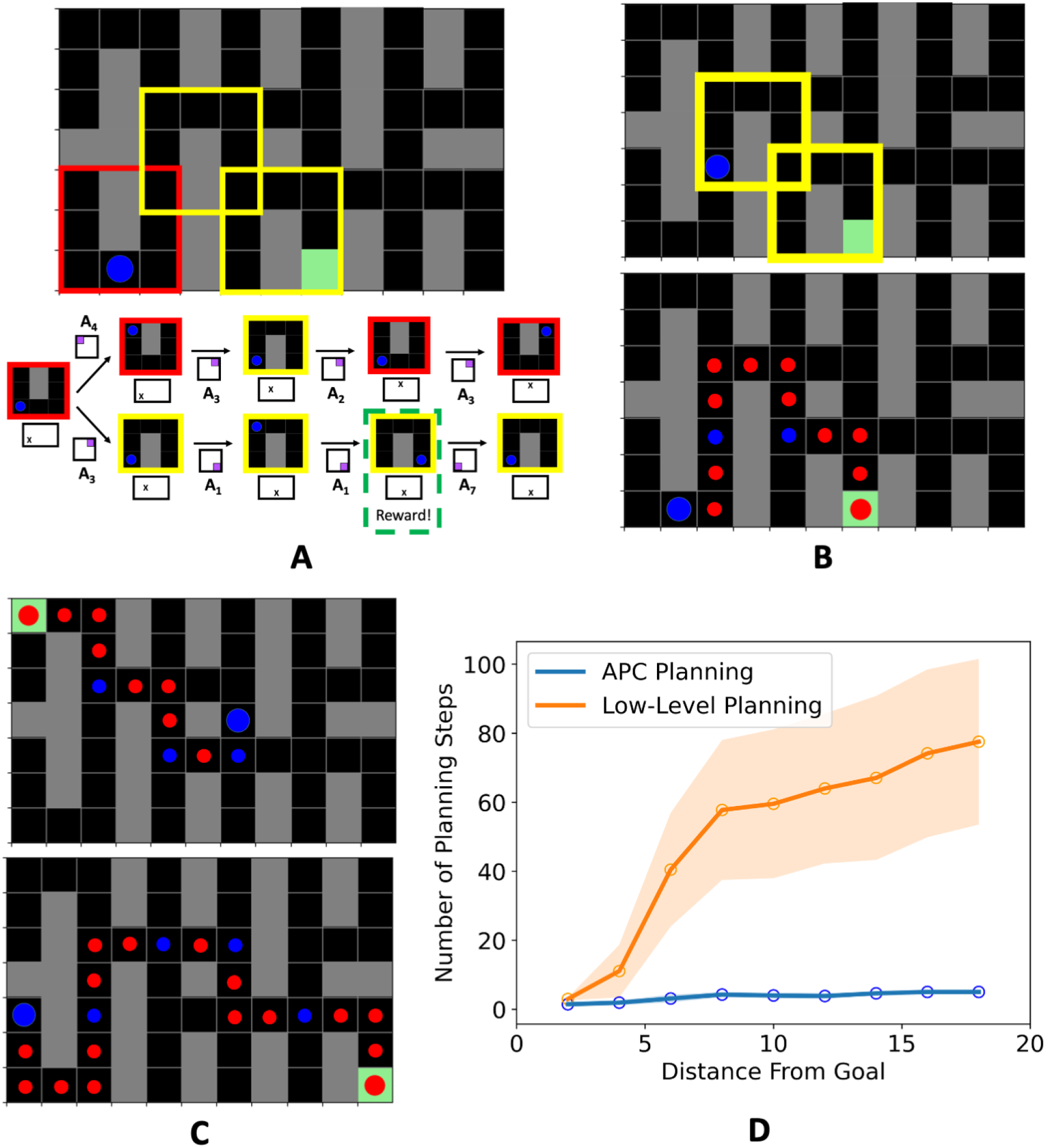
Hierarchical Planning. (A) To navigate to the green goal location from the blue start location, the APC model uses its learned high-level state network to sample *K* high-level state-action sequences (*K* = 2 here, shown bifurcating from the initial state). In each sequence, the high-level state is depicted by a predicted room image (red or yellow outlined image) and its location (marked by an “X” in the global frame below the image). High-level action is depicted by its associated goal location (purple) in a square local frame. (B) Given the sampled sequences, the model picks the sequence with highest total reward, executes this sequence’s first (high-level) action to reach the blue location shown in the top panel, and repeats to reach the goal location with 3 high-level actions (bottom panel in (B)). Small red dot: intermediate location; small blue dot: intermediate goal. (C) shows the same APC model solving two more navigation problems involving new start and goal locations using hierarchical planning. (D) High-level planning by the APC model versus low-level heuristic planning using primitive actions (see text for details). Average number of planning steps for the low-level planner quickly increases as distance to goal increases compared to high-level planning by the APC model. (Adapted from [10]).

Figure 9D illustrates the efficacy of the APC model’s high-level planning compared to lower-level planning (MPC using random sequences of 4 primitive actions; Euclidean distance heuristic; see [10] for details): as expected, the average number of planning steps to reach the goal increases dramatically for larger distances from the goal for the lower-level planner compared to high-level planning by the APC model.

## 5 Cortical Predictive Coding, Hippocampal Binding and Episodic Memory

Before concluding, we note that the state and action networks at different levels of the APC hierarchy learn generic “basis functions” in their synaptic weights for representing states and actions. The basis functions emerge as result of learning from interactions with the environment. The simplest example of these basis functions are at the lowest level of the hierarchy in our APC example for vision. When a sparseness constraint is imposed on the state vectors [49], the basis functions that are learned from natural videos comprise of oriented Gabor filters coding for edges/bars at different orientations (for more details, see results in [48] from a version of the APC model without actions (Dynamic Predictive Coding)). In this case, the state vector 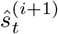 is a specific activation pattern of the state-estimating neurons representing a specific image patch as a combination of the learned basis Gabor filters.

Generalizing this idea to the APC hierarchical network, it can be seen that at the highest level *N* of the network, we have specific activation patterns *ŝ*^(*N*)^ and *â*^(*N*)^ of state-estimating neurons and abstract action/goal-encoding neurons together representing an entire sequence of specific states and actions corresponding to the current episode of interaction with the environment.

By connecting this highest level of the APC network to a hippocampus-like associative memory, one can store the current activation patterns *ŝ*^(*N*)^ and *â*^(*N*)^ as an episodic memory vector *m* (Figure 10A). This episodic memory can later be retrieved when given an internal or external cue, e.g., a partial input that is the beginning of the episodic sequence. Figure 10B (from [48]) provides an example of such a recall by a two-level memory-augmented dynamic predictive coding model (APC without actions) which was shown 5 episodes of a sequence depicting the digit “5” moving from left to right (top panel in Figure 10B). By storing the sequence information as a vector *m* in its associative memory, the model was later able to retrieve the entire sequence given only the starting image as a cue (Figure 10B (lower panels)): this is reflected in the similarity between the activation pattern of the network’s lower-level neurons during recall (“Start” condition) and the activation pattern observed during training (“Conditioning”).

**Figure 10:**
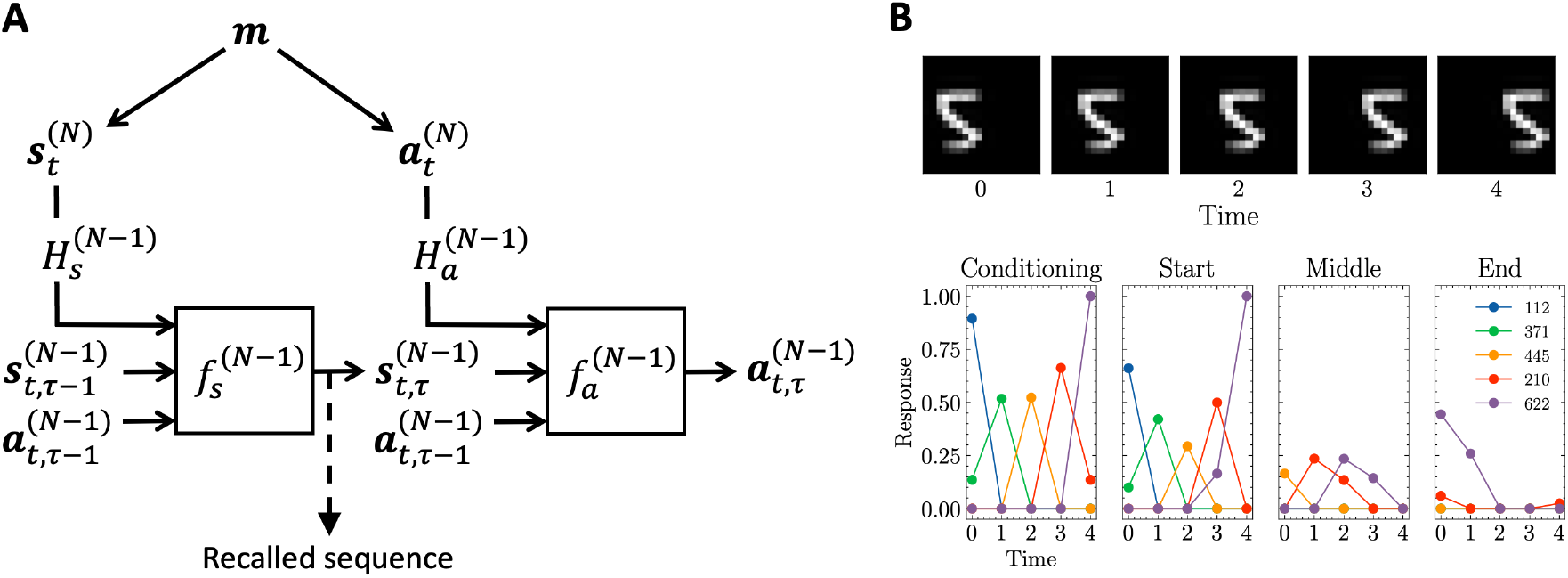
Hippocampal Binding of Cortical Representations and Episodic Memory. (A) depicts a generative model that connects an associative memory (emulating the hippocampus) to the highest level of a hierarchical APC model (emulating the cortex). The associative memory stores a memory vector *m* that encodes the current highest-level state and action vectors *ŝ*^(*N*)^ and *â*^(*N*)^ as an episodic memory. Later activation of the memory vector *m* by an internal or external cue recalls the entire sequence of states and actions in the memorized episode as activations cascade down the cortical hierarchy. (B) An example of episodic recall in a two-level memory-augmented model. The model was shown 5 episodes of the image sequence in the top panel. The bottom panel shows the responses of five most active lower-level model neurons at each time step when presented with the training image sequence (“Conditioning”) and when presented with only the “Start” frame (image from time step 0), only the “Middle” frame (from time step 2) and only the “End” frame (from time step 4). Episodic recall from memory is indicated by the similarity in activation patterns between “Conditioning” and “Start.” ((B) adapted from [48]).

More importantly, the cortical representation at the highest levels can be regarded as a factored representation of the world, involving state representations from multiple modalities (vision, audition, somatosensation etc.) and actions. The convergence of these different state and action vectors into an associative memory at the top of the hierarchy allows the model to perform *binding*: these different multimodal representations are bound into a single temporary memory representation *m*. This representation is fed back to the cortical network as depicted in the generative model in Figure 10A, allowing the fused multimodal information in memory to influence the different cortical areas down the hierarchy to the lowest levels.

The above ideas are consistent with the notion of binding in AI, particularly in neurocompositional representations, where “fillers” (specific instances) are bound to semantic “roles” and allow modeling of lingistic, logical and symbolic concepts [35]. We expect the benefits of such a representation, which include fast transfer of knowledge and zero shot learning due to compositional generalization, to also accrue to the memory-augmented APC model. Such a model shares similarities with the Tolman-Eichenbaum Machine (TEM) model of interactions between the hippocampus and the entorhinal cortex [63].

In summary, the APC model suggests that the cortex encodes generic semantic knowledge about the world within state and action networks that implement nested reference frames. Any particular instantiation of this knowledge invoked by, for example, an interaction with a person or an object, is stored temporarily as an episodic memory vector *m* in the hippocampus. This instantiation could be used for reasoning about the current situation or for planning, and if deemed important, could be consolidated within the cortex by updating cortical networks via replay during inactivity or sleep.

## 6 Discussion

This article explored a sensory-motor theory of cortical function based on active predictive coding (APC). The theory proposes that (a) each cortical column implements both a state transition network for state prediction and a policy network for action/goal prediction (thereby defining a reference frame [7, 27]), and (b) higher-level neurons representing more abstract states and actions modulate the lower-level state and action networks via top-down modulatory feedback to change the functions they are computing, leading to nested reference frames and hierarchical representations of objects, states and actions. A neuroanatomical mapping of the APC model to cortical laminar structure was suggested in Section 2.4, although it should be noted that this is only one of several possible mappings and key elements of this mapping remain to be tested.

### 6.1 Diverse Capabilities of the APC Model

The APC model confers on the cortex a diverse range of capabilities using the same basic architecture, lending support to the hypothesis [5–7, 64, 65] of a common computational principle operating across the neocortex:

- **Parsing images and learning part-whole hierarchies**: Eye movements can be used to parse images and learn hierarchical representations of parts and sub-parts of objects (Section 4.1.1);
- **Invariant perception**: Learned representations of objects and sequences can be transformed by the generative model to match current inputs to remain invariant to different types of transformations (translations in Section 4.1.1, other transformations such as rotations and scaling in [12]);
- **Perceptual stability**: Inference in the APC model naturally leads to integration of information across actions such as eye movements, leading to perceptual stability (Section 4.1.1);
- **Compositionality and fast transfer of knowledge**: The APC architecture allows compositional representations to be learned, allowing the model to compose and generate new objects and sequences made up of novel combinations parts and sub-parts, allowing fast generalization and recognition of novel inputs (Sections 4.1.1, 4.2 and 5);
- **Efficient planning**: Hierarchical state networks in the APC model can be used to solve tasks such as navigating in a large building efficiently by planning using hierarchical actions (Section 4.2);
- **Habit formation**: Successful plans can be used to learn new policies (“habits”); alternately, the APC model also allows policies to be learned using hierarchical reinforcement learning (Section 4.1.1);
- **Reference frames and temporal hierarchies**: The APC model provides a neural implementation of the concept of nested reference frames [7] and shows how joint temporal hierarchies of states and actions can be learned;
- **Prediction and postdiction**: Since the model maintains a temporally stable state representation at the highest level that encodes entire sequences, the update of this representation during prediction error minimization leads to an explanation for both predictive and postdictive phenomena in perception such as the flash-lag illusion and the color-phi effect (see [48] for details);
- **Generating “schemas” or “programs” for solving novel tasks**: The APC model suggests a neural mechanism (via top-down inputs/gain modulation) for generating new sensory-motor “programs” on-the-fly to solve novel tasks (Section 3.4);
- **Binding and episodic memories of perception-action sequences**: When coupled with an associative memory emulating the role of the hippocampus, the model binds multimodal cortical activations at the highest level into an episodic memory (sequence of multimodal states and actions), allowing activity recall and cortical consolidation at a later time, and promoting fast generalization and learning (Section 5);
- **Language and symbolic representations**: The state-action representations in the APC model can be made categorical [57], opening the door to representing symbols, learning grammars and understanding and producing language. The ability of the APC model to perform binding with the current task’s cortical representation is similar to the “filler:role” mechanism used in neurocompositonal computing [35] for solving linguistic and symbolic tasks;
- **Learning abstract concepts**: The same sensory-motor architecture used for perception and planning can also be used to model abstract concepts such as family trees by using abstract states to represent parents and children, and abstract actions (up, down etc.) to traverse and define the tree; results along these lines were obtained using the TEM model [63] where a recurrent neural network (similar to our state transition network) was used in conjunction with an associative memory to learn the structure of family trees from examples.

### 6.2 Related Models

The APC model shares broad similarities with other models of cortical function. The idea that the cortex relies on predictions and performs inference over a hierarchy of time scales is common to many models of cortical function [1, 66–71], going back to the seminal early work of MacKay [72] and Albus [73]. The goal of putting action on an equal footing with perception in terms of Bayesian inference and prediction error minimization is in keeping with the theories of active inference and free energy minimization proposed by Friston and colleagues [50, 74]. Compositionality and the representation of sensory-motor information in cortical columns are also central tenets of the “thousand brains” theory proposed by Hawkins and colleagues [7, 27, 28]. Their model differs from the APC model in postulating the existence of grid cells in all cortical areas, not leveraging hierarchical state-action representations, and not invoking policy functions for computing actions.

The close interaction between state-estimation networks and action-computing networks in the APC model is consistent with theories of optimal motor control [75]. However, based on recent evidence pointing to motor outputs from Layer 5 in essentially all cortical areas [4, 13–15, 17], the APC model proposes that all cortical columns include both state-estimation and policy components. Even primary motor cortex (M1), which is often cited as an example of a cortical area missing the sensory input Layer 4, receives sensory information from other cortical and subcortical areas [76–78], and can therefore predict and estimate state (e.g., proprioceptive state) and compute actions via the circuits in the superficial and deep layers of M1. In fact, the APC model’s interpretation of Layer 5 outputs as abstract actions or goal vectors for spinal circuits is consistent with a previously proposed active inference model of M1 [33].

Finally, the idea that the policy network outputs serve as a prior for selecting actions during planning provides a potential neural implementation of Gibson’s model of perception based on *affordances* [79]. An object, when estimated by higher-level state-estimating neurons, generates a prior distribution over suitable actions for that object via the policy network in the APC model (see Section 3.3.2). This prior affinity for certain actions for an object is learned by the hierarchical cortical state-action networks during the course of solving many tasks involving similar types of objects.

### 6.3 Model Predictions and Open Questions

The first major prediction of the APC model is that every cortical column computes an *abstract action* or *“goal”* that is conveyed to subcortical motor centers, which may or may not choose to act upon this output. The actual action that was executed is then conveyed back to all cortical areas via the thalamus to update the cortical state and action representations. The precise mechanism that mediates the final action selection in subcortical motor centers based on cortical outputs remains an open question.

A second prediction is that cortical areas interact with each other via top-down modulation targeting apical dendrites of cortical pyramidal neurons – the computational role of such modulation is to *change the function* being computed by the lower level’s state-transition and action/policy networks based on a higher-level’s state and action vectors (Section 3.2.1). Such an arrangement predicts that representations at higher levels encode information over longer time scales than lower levels [8, 9, 48]. The mechanisms underlying the modulatory interactions between higher-order and lower-order cortical areas across these time scales, and the role of alpha, beta, theta and gamma oscillations in such interactions are not yet clear.

While there is emerging neurophysiological and neuroanatomical evidence that lend some support to the APC model’s predictions [13, 29–33, 38], there is much that remains to be tested. We hope that the APC theory inspires new experiments that address the core question that motivated the theory: how does the neocortex learn and use its sensory-motor internal model of the world?

## Acknowledgments

The author would like to thank members of the “Predictive Coding Task Force” in the Neural Systems Laboratory at the University of Washington, Ares Fisher, Dimitrios Gklezakos, Preston Jiang, Prashant Rangarajan and Vishwas Sathish, for many discussions and the collaborative work cited in the text. The author would also like to thank Saghar Mirbagheri, Nick Steinmetz, Giedrius Burachas, Eb Fetz, Adrienne Fairhall and Eric Shea-Brown for discussions and feedback on ideas related to predictive coding. This research is currently supported by National Science Foundation (NSF) EFRI Grant no. 2223495, a Weill Neurohub Investigator grant, and a “Frameworks” grant from the Templeton World Charity Foundation, and is based partly upon work supported previously by National Science Foundation (NSF) Grant no. EEC-1028725 and the Defense Advanced Research Projects Agency (DARPA) under Contract no. HR001120C0021. The opinions expressed in this publication are those of the author and do not necessarily reflect the views of the funders.

## Footnotes

1 A similar biasing strategy for planning actions may also be used for imitation-based learning, where the observed actions from an expert solving a task can be used to bias one’s own action search during planning to quickly solve the same task (see, e.g., [58]).

